# *Wolbachia* enhances the survival of *Drosophila* infected with fungal pathogens

**DOI:** 10.1101/2023.09.30.560320

**Authors:** Jessamyn I. Perlmutter, Aylar Atadurdyyeva, Margaret E. Schedl, Robert L. Unckless

## Abstract

*Wolbachia* bacteria of arthropods are at the forefront of basic and translational research on multipartite host-symbiont-pathogen interactions. These microbes are vertically inherited from mother to offspring via the cytoplasm. They are the most widespread endosymbionts on the planet due to their infamous ability to manipulate the reproduction of their hosts to spread themselves in a population, and to provide a variety of fitness benefits to their hosts. Importantly, some strains of *Wolbachia* can inhibit viral pathogenesis within and between arthropod hosts. Mosquitoes carrying the *w*Mel *Wolbachia* strain of *Drosophila melanogaster* have a greatly reduced capacity to spread viruses like dengue and Zika to humans. Therefore, *Wolbachia* are the basis of several global vector control initiatives. While significant research efforts have focused on viruses, relatively little attention has been given to *Wolbachia*-fungal interactions despite the ubiquity of fungal entomopathogens in nature. Here, we demonstrate that *Wolbachia* increase the longevity of their *Drosophila melanogaster* hosts when challenged with a spectrum of yeast and filamentous fungal pathogens. We find that this pattern can vary based on host genotype, sex, and fungal species. Further, *Wolbachia* correlates with higher fertility and reduced pathogen titers during initial fungal infection, indicating a significant fitness benefit. This study demonstrates *Wolbachia*’s role in diverse fungal pathogen interactions and determines that the phenotype is broad, but with several variables that influence both the presence and strength of the phenotype. These results enhance our knowledge of the strategies *Wolbachia* uses that likely contribute to such a high global symbiont prevalence.

**Importance:** *Wolbachia* bacteria of arthropods are at the forefront of global initiatives to fight arthropod-borne viruses. Despite great success in using the symbiont to fight viruses, little research has focused on *Wolbachia*-fungal interactions. Here, we find that *Wolbachia* of *Drosophila melanogaster*, the same strain widely used in antiviral initiatives, can also increase the longevity of flies systemically infected with a panel of yeast and filamentous fungal pathogens. The symbiont also partially increases host fertility and reduces fungal titers during early infection, indicating a significant fitness benefit. This represents a major step forward in *Wolbachia* research since its pathogen blocking abilities can now be extended to a broad diversity of another major branch of microbial life. This discovery may inform basic research on pathogen blocking and has potential translational applications in areas including biocontrol in agriculture.

## Introduction

Microbe-host symbioses are ubiquitous in nature and exhibit a broad range of relationships from facultative parasitism to obligate mutualism^1,2^. Microbial symbionts of arthropods in particular exhibit a striking array of phenotypes in their hosts^2^, ranging from provision of nutrients^3^ to protection from parasitoids^4^ to death of the host’s offspring^5^. One microbial symbiont, *Wolbachia pipientis*, is an exemplary case of a microbe with diverse symbiont-host interactions. *Wolbachia* are obligate intracellular bacteria found in germline and somatic tissues of diverse arthropods and are almost exclusively inherited vertically through the cytoplasm of infected mothers^6^. They are found in an estimated 40-52% of all arthropod species on Earth^7,8^, making them the most widespread endosymbiont and “the world’s greatest pandemic”^9,10^. There is such genetic diversity that there are 18 recognized *Wolbachia* supergroups^11–13^. Some can act as “reproductive parasites” that manipulate host reproduction to facilitate their spread by enhancing the relative fitness of infected female transmitters^14^. Others are obligate mutualists necessary for host oogenesis or early development^15^. Depending on context, *Wolbachia* can use their diverse genetic toolkit to engage in a variety of interactions with their hosts. These interactions have had immense impacts on both basic and applied research in many fields, including utility in fighting human diseases vectored or caused by insects and nematodes and to an understanding of the role of symbionts in shaping host evolutionary processes^6,16–18^.

*Wolbachia*’s employment of such diverse host interactions has been critical to its global success, however, these phenotypes do not fully explain how widespread *Wolbachia* is. Indeed, while some strains are reproductive manipulators (enhancing the fitness of the infected matriline)^5,10,19–21^ or obligate mutualists (enhancing the fitness of all hosts)^12,22–24^, but many are not, even among organisms that have been phenotypically assessed^25^. Some strains also exhibit no reproductive parasitism in and provide no currently known fitness benefit^26,27^. Further, those that are reproductive manipulators can vary both in the effect size of their phenotype (either weak or strong induction^28–31^) and in their frequency in the population (high or low^32–35^). Even when reproductive phenotypes or benefits are known, they are often context-dependent and vary based on factors such as temperature^36–39^, symbiont density^40,41^, or host genetic background^42^. Further, in the wild, vertical transmission fidelity of *Wolbachia* is not 100%^27,43,44^, making the basis of the symbiont’s maintenance in populations even less clear. For many years, a question of significant focus in the field has been how it is that *Wolbachia* is so widespread^45^, particularly given the fact that we have not identified a clear host fitness benefit of the symbiont for all strains or contexts. Research over the years has identified some contributing factors such as nutritional contributions of the symbiont to the host^46,47^, as well as rescue of host deficiencies like mutations in the key sex development regulator *sex-lethal*^48,49^ and germline stem cell self-renewal and differentiation deficiencies^50^. Yet these contributing factors do not fully answer the question, and other factors must be involved.

One such crucial and somewhat common beneficial *Wolbachia*-host interaction was discovered through work on an early theory that *Wolbachia*’s prevalence could be based on an ability to inhibit pathogens, thereby conferring a significant fitness benefit to the host^51–53^. The rationale was based partially on the observation that facultative infection (as opposed to obligate mutualism) is relatively common with *Wolbachia* infections, but with few accompanying known benefits to explain their frequency. It was also partially based on an observation that *Wolbachia* infection correlated with host resistance to infection with the common *Drosophila* C virus (DCV)^51^. Two foundational early studies on this topic demonstrated that *Drosophila melanogaster* flies with their native *Wolbachia* strain exhibit greater longevity on the order of days to weeks of increased life when infected with several common arthropod RNA viruses^51,54^. This coincides with reduced viral load in *Wolbachia*-viral co-infection, which increases host fitness and survival likelihood though reduced pathogen burden. These and latter studies also demonstrated that the phenotype could be induced by some additional *Wolbachia* strains or in additional host genetic backgrounds or species, but that the effect was largely restricted to RNA viruses (not DNA viruses)^55^. Finally, and crucially, some *Wolbachia* strains are also able to inhibit the transmission of viral (and some other) pathogens to new host individuals, including pathogens spread by mosquitoes to humans^51,54,56,57^. This ability of the symbiont to protect its host from viruses is considered a major factor contributing to *Wolbachia*’s success.

Virus pathogen blocking has therefore become an eminent area of *Wolbachia* research not only for its broad applicability across the symbiont genus and importance to basic biology, but also for its translational potential. For example, *Aedes aegypti* mosquitoes and other common human disease vectors exhibit significantly reduced capacities to transmit parasites like malaria^57^ or viruses like Zika^56^, dengue^58,59^, yellow fever^60^, or chikungunya^61^ to humans when they carry certain strains of *Wolbachia*. This feature has made *Wolbachia* central to global efforts to reduce disease through groups like MosquitoMate^62^ and the World Mosquito Program^63^. These programs rear *Wolbachia*-positive mosquitoes on a massive scale and release mosquitoes into the wild. One strategy is to release infected females that then outcompete local *Wolbachia*-negative counterparts and replace them with a disease-resistant population. Collaborative efforts through this program across four continents have resulted in stable, wild *Wolbachia*-positive populations in many locations and significant reductions in disease^58,64^. Arthropod vector-borne diseases are responsible for millions of illnesses, deaths, and contribute to significant inequality around the world^65^, and the use of *Wolbachia*-positive mosquitoes is one of our most promising solutions^66–68^.

In contrast with all of this progress on viruses, comparatively little research has been done on *Wolbachia* interactions with non-viral pathogens^57,69^. This is despite the extraordinary genetic and phenotypic diversity of *Wolbachia* symbioses that indicate the likelihood of broader protective abilities. Early theory predicted that pathogen protection could increase the relative fitness of hosts with *Wolbachia* compared to those without, contributing to maintenance and spread of the symbiont^32^, and this was one of the original bases for investigations into viral pathogen blocking, and could apply to many other types of pathogens too^51,54^. However, one particular gap in the research is the potential for *Wolbachia* to inhibit fungal pathogens. Fungal pathogens of arthropods are common in the wild^70^, yet few studies have investigated the interactions between *Wolbachia*, hosts, and fungal pathogens, and the studies that do present different results. One early study showed no effect of *w*Ri *Wolbachia* strain infection on survival from topical cuticle infection of the common insect fungal pathogen, *Beauveria bassiana*, in *D. simulans* male flies^71^. Another reported higher survival of *D. melanogaster* female flies with their native *w*Mel *Wolbachia* symbiont after immersion in a suspension of *B. bassiana*^72^.

Conversely, a third study on infection of female spider mites in topical contact with *B. bassiana* or *Metarhizium* fungal pathogens indicated that *Wolbachia* may actually increase mortality of the host with fungal infection^73^. A fourth investigated the effect of *Wolbachia* on injection with two *Beauveria* pathogens on *Aedes albopictus* and *Culex pipiens* mosquitoes^74^. This study found no enhancement in host survival with the symbiont, but reported some putative differences in host immune gene expression and reduced fungal load in some contexts. Finally, a recent study indicates that the *w*Pni strain of *Pentalonia* aphids may result in increased survival of hosts infected topically with the specialized fungal pathogen, *Pandora neoaphidis*^75^. Thus, there have been several investigations, with some prior reports indicating that *Wolbachia* may interact with fungal pathogens in some contexts.

Despite this research, the question of *Wolbachia*’s ability to interact with fungal pathogens on a larger scale remains unanswered. It is unclear how broad the fungal blocking ability is in terms of host, symbiont, and pathogen factors, and if the phenotype is likely to be common or not. This difficulty is because the studies draw different conclusions from different contexts. These prior reports have used different host species, host sexes, *Wolbachia* strains, pathogen species, pathogen concentrations, routes of pathogen infection, and been measured by different host fitness and health assays or conducted over different lengths of time^71–75^. These factors make it difficult to compare across studies, as there are multiple variables between any two publications. Further, due to the small number of studies, limited parameters have been tested thus far. Thus, the breadth of *Wolbachia*-fungal interactions is unclear, as comparison between studies is difficult and there is limited published data.

To begin to fill this gap in knowledge, we conducted a series of systemic fungal infection assays using *D. melanogaster* flies with the *w*Mel *Wolbachia* symbiont in the context of several host and pathogen variables. Notably, *w*Mel is the initial strain that was reported to inhibit viruses and mosquitoes transinfected with this symbiont strain are the basis of many of the global vector control initiatives^51,54,58^. This approach addresses several outstanding research questions in this area: (i) can *Wolbachia* inhibition of fungal pathogenesis be confirmed when tested in various contexts, (ii) how broad is this protective phenotype within one *Wolbachia* strain, and (iii) do factors such as fungal pathogen species, fungal pathogen types (filamentous vs yeast), host sex, and host genetic background contribute to the *Wolbachia*-fungal pathogen interaction. Here we report that *Wolbachia* is indeed capable of significantly increasing the longevity and reproductive fitness of flies infected with a wide variety of fungal pathogens, and the phenotype is influenced by several host and pathogen factors.

## Results

### *Wolbachia’s* association with an increase in longevity of flies infected with filamentous fungi is dependent on genetic background and host sex

To test the breadth and ability of *Wolbachia* to inhibit fungal pathogenesis in flies, a series of systemic infection assays were conducted. Experiments were performed with two different *Drosophila melanogaster* host background lines infected with their native *w*Mel *Wolbachia*. The host strains themselves have diverse origins: the *w*^*1118*^ line was collected in California, USA and was reported in 1985^76^, and the *w^k^*line was collected in 1960 in Karsnäs, Sweden^77^. Different collection origins together with Illumina sequencing showing a high number of SNPs between the *D. melanogaster* lines indicate the lines represent genetically diverse host backgrounds. Each strain has its own natural *Wolbachia* along with genetically identical counterpart strains that were previously treated with antibiotics to remove the symbiont. Thus, we tested four strains total: *w*^*1118*^ with *Wolbachia*, *w*^*1118*^ without *Wolbachia*, *w^k^* with *Wolbachia*, and *w^k^* without *Wolbachia*. Whole genome sequencing of the *Wolbachia* symbionts of each strain indicates that they are highly similar despite disparate origins, with only a single divergent SNP across the entire genome. This SNP is a silent (synonymous) polymorphism in a membrane transporter of the major facilitator superfamily, which transports small solutes^78^. Thus, the vast majority of genetic differences between strains can be attributed to the host, and most phenotypic differences are therefore likely due to the host as well.

To determine if *Wolbachia* can increase the longevity of flies infected with fungi as hypothesized, systemic infections were performed with both sexes of all four strains against a variety of pathogens. We started with several *Aspergillus* and *Fusarium* filamentous fungal species that infect both arthropods and humans: *Aspergillus fumigatus*, *Aspergillus flavus*, *Fusarium oxysporum*, and *Fusarium graminaerum* (Figure 1). Survival was scored daily for three weeks, as differences in survival were broadly apparent across treatment groups for most pathogens by this point. The data revealed several key results. First, *Wolbachia* was associated with significantly greater survival across the trial period in many contexts. In the *w^k^* background, *Wolbachia*-positive flies had higher survival for all pathogens except *Fusarium oxysporum*, which was only significant when comparing within just males (Figure 1). Second, genetic backgrounds played a significant role in the infection outcomes. Indeed, *Wolbachia* was not a significant predictor of increased longevity for any of the pathogens in the *w*^*1118*^ host background, except when considering sex (Figure S1). Third, sex is repeatedly a significant factor in survival outcomes for some pathogens. Males alone had a significant increase in longevity for *Aspergillus fumigatus* and *Fusarium oxysporum* for both genetic backgrounds (Figures 1a,c & S1a,c), with a statistically significant *Wolbachia* x sex interaction for *A. fumigatus in* the *w^k^* background and *Fusarium oxysporum* in the *w*^*1118*^ background (Figures 1a, S1c). Fourth, the host strains had generally different overall susceptibilities to fungal infection, with *w^k^* generally having lower survival than *w*^*1118*^ in both *Wolbachia*-positive and -negative contexts (Figures 1 & S1, mean 51.1% death for all pathogen infections combined in the *w*^*1118*^ background by day 21, 60.4% death in the *w^k^* background). In particular, there is a significant *Wolbachia* x genotype interaction for *Aspergillus flavus* (*p=0.043, Table S1).

**Figure 1.**
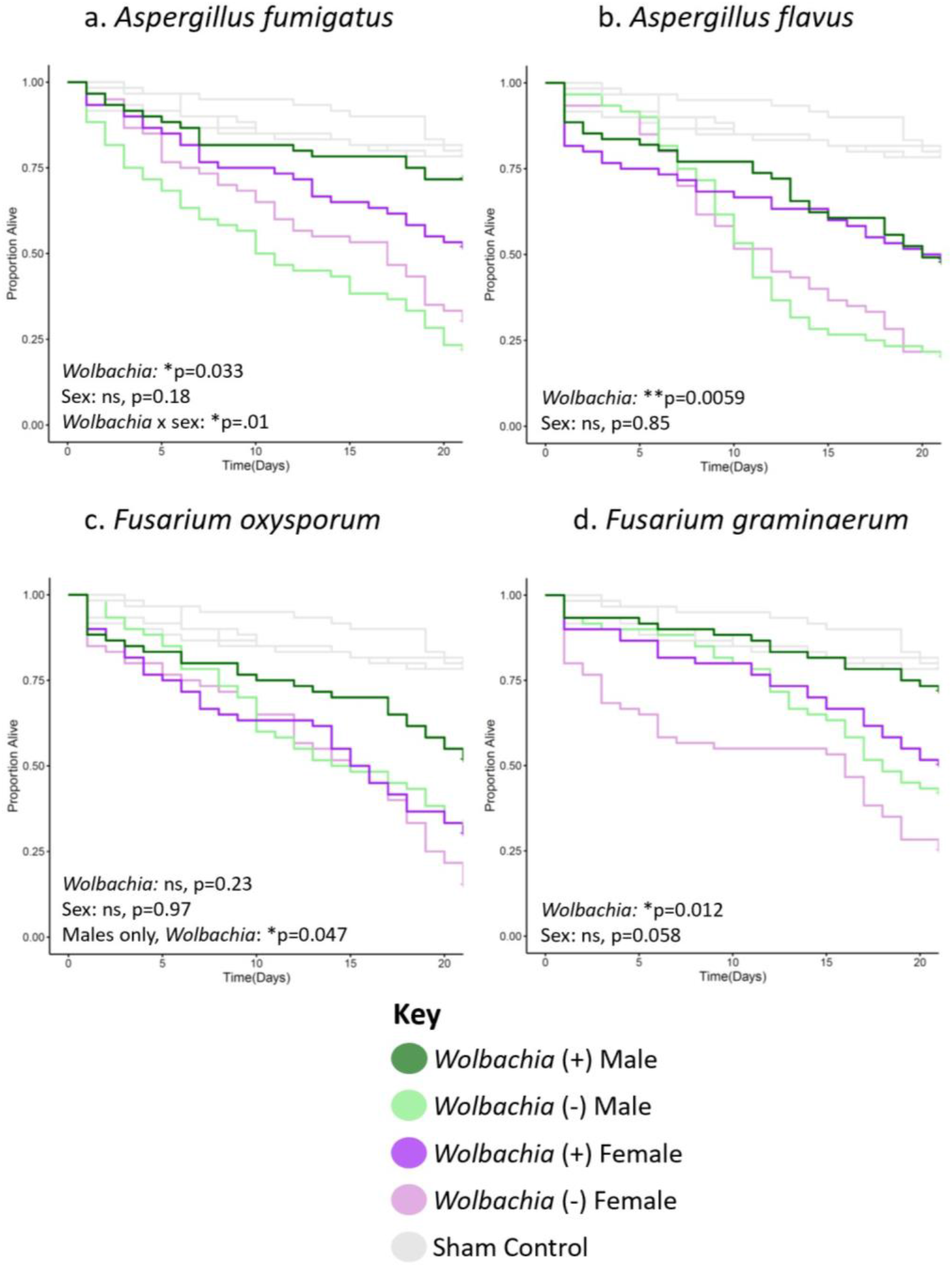
*Wolbachia* increases the longevity of flies of the *w^k^* background line infected with several filamentous fungal pathogens. Flies of each given background and sex were systemically infected with the indicated pathogen. Infections were performed with either (a) *Aspergillus fumigatus*, (b) *Aspergillus flavus*, (c) *Fusarium oxysporum*, or (d) *Fusarium graminaerum*. Infections of all groups were performed side-by-side, along with those of the *w*^*1118*^ background line (Figure S1), with at least two blocks of infections performed on different days. Each line represents a total of 60 flies. Sham controls were performed with sterile 20% glycerol. Full statistics, available in Table S1, were done with a Cox mixed effects model. Controls are the same in all panels and in Figure 2a because they were performed concurrently in the same background.

### *Wolbachia* can increase the longevity of flies infected with filamentous fungal entomopathogens

To determine if *Wolbachia* could also increase longevity of flies infected with common filamentous fungal insect pathogens (entomopathogens), we performed systemic infections with *Beauveria bassiana*, *Metarhizium anisopliae*, *Clonostachys rosea*, and *Trichoderma atroviride*. *Beauveria* and *Metarhizium* in particular are ubiquitous insect pathogens and are the subject of extensive research in biocontrol of pests in particular^79^, while *Clonostachys* and *Trichoderma* are also globally widespread and have received recent attention in biocontrol as well^80–82^. The latter two were collected from mosquitoes, and are thus of potential relevance to mosquito biology (Table S2). Similar to the results of the pathogens in Figures 1 & S1, *Wolbachia* increased longevity in many, but not all fungal infection contexts (Figures 2 & S2). Namely, *Wolbachia* significantly increased longevity for *Beauveria bassiana* and *Clonostachys rosea* in the *w^k^*background (Figure 2a,c), and *Beauveria bassiana* and *Metarhizium anisopliae* in the *w*^*1118*^ background (Figure S2a,b). Thus, there is some positive longevity effect of the symbiont in either background, not just *w^k^*, but the effect depends on the pathogen. Further, sex was also a factor with a significant effect for *Beauveria bassiana* and *Metarhizium anispoliae* in the *w^k^* background (Figure 2a,b) and *Metarhizium anisopliae* and *Trichoderma atroviride* in the *w*^*1118*^ background (Figure S2b,d). Additionally, as with previous infections, *w^k^* was broadly more susceptible to infection as flies generally died earlier and at higher rates than their *w*^*1118*^ counterparts (Figures 2 & S2, mean 70.3% death for all entomopathogen infections combined in the *w*^*1118*^ background by day 21, 85.8% death in the *w^k^* background).

**Figure 2.**
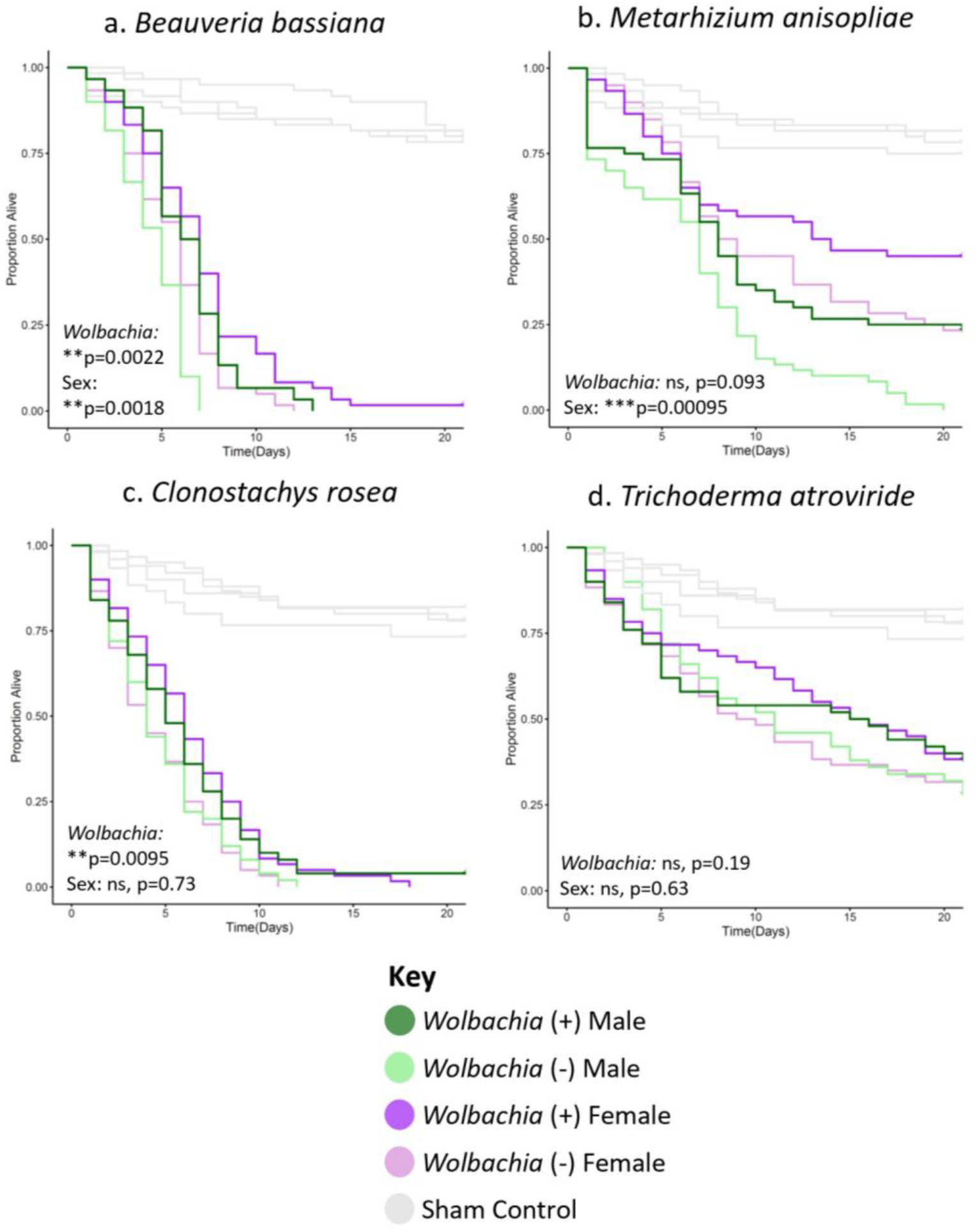
*Wolbachia* increases the longevity of flies of the *w^k^* background line infected with certain filamentous fungal entomopathogens. Flies of each given background and sex were systemically infected with the indicated pathogen. Infections were performed with either (a) *Beauveria bassiana*, (b) *Metarhizium anisopliae*, (c) *Clonostachys rosea*, or (d) *Trichoderma atroviride*. Infections of all groups were performed side-by-side, along with those of the *w*^*1118*^ background line (Figure S2), with at least two blocks of infections performed on different days. Each line represents a total of 60 flies. Sham controls were performed with sterile 20% glycerol. Full statistics, available in Table S1, were done with a Cox mixed effects model. Controls for panel 2a are the same for Figure 1, and the panels in 2b-d are the same because they were performed concurrently in the same background.

### *Wolbachia* can increase the longevity of flies infected with yeasts

To test if *Wolbachia* could also increase the longevity of flies infected with yeast, we performed systemic infections using *Candida auris, Candida glabrata*, and *Galactomyces pseudocandidus.* For *Candida* pathogens, *Wolbachia* significantly increased longevity of *w^k^*background flies. In contrast, *Wolbachia* did not significantly increase longevity for any of the yeast pathogens in the *w*^*1118*^ background. Further, sex was not a significant factor in any of the yeast infections for either background. However, flies of the *w^k^* background again were more broadly susceptible to infection based on higher overall mortality (mean 40% death for all yeast infections combined in the *w*^*1118*^ background by day 21, 58.3% death in the *w^k^* background).

### *Wolbachia* can partially rescue female fertility reduction after infection

To assess whether *Wolbachia* impacts fitness of hosts early in fungal infection, female flies were systemically infected with *B. bassiana* because *Wolbachia* significantly increased longevity for all treatment groups with this pathogen (Figures 2a, S2a). Egg laying and egg hatching rates were quantified for the first 3 days post infection for flies with either the infection or a sham control (Figures 4, S4). Although both *Wolbachia*-positive and *Wolbachia*-negative flies laid similar numbers of eggs in the *w^k^* background without treatment, and although the overall egg-laying was lower in *B. bassiana*-infected flies, *Wolbachia* significantly increased egg-laying with fungal infection (Figure 4). This was also true in the *w*^*1118*^ background (Figure S4). In contrast, the percentage of eggs hatched was not greatly impacted by either *Wolbachia* or fungal infection in either background (Figures 4b, S4b).

### *Wolbachia* associates with reduced fungal titer after infection

To determine if enhanced longevity is likely based on killing or reduction of pathogen (immune resistance) vs tolerance and maintenance of the pathogen (immune tolerance), and to determine if reproductive benefits with fungal infection in Figures 4 & S4 can be attributed to reduced pathogen load, we measured fungal and *Wolbachia* titers over time in *B. bassiana*-infected females (Figure 5). We measured over the first 24 h because this is before flies begin to die and many essential early host molecular responses to pathogen infection begin by this timepoint during infection^83,84^. We find that *Wolbachia* titer stays constant over the 24 h period (Figure 5a) and that pathogen load is not significantly different between lines immediately post-infection (Figure 5b). Thus, both *Wolbachia*-positive and -negative flies are receiving similar starting amounts of pathogen. However, by 24 h post-infection, we see that pathogen load is reduced in the *Wolbachia*-positive flies compared to those without *Wolbachia*. This trend holds true in the *w*^*1118*^ background as well.

## Discussion

In the 15 years since the discovery of *Wolbachia*-based virus inhibition, there has been significant research into the mechanism and translational applications of the phenotype^51,54,55,64^. However, comparatively little attention has been given to the potential for *Wolbachia* to interact with other types of pathogens, including fungi. Prior research gave contrasting results either suggesting there was a *Wolbachia*-fungal infection interaction^72,75^ or not^71,73,74^. However, these previous studies were performed in different contexts with many different variables between them. Thus, the breadth of *Wolbachia*’s ability to interact with fungal pathogens as well as identification of factors that influence the putative phenotype have remained unclear. Given the likely importance of fungal interactions to the basic biology of *Wolbachia* and potential applications in areas like agriculture, these are important research topics to address. For example, the large field trials that release *Wolbachia*-positive mosquitoes to combat arthropod-transmitted viruses rely on *Wolbachia*’s reproductive manipulations of the host to help spread itself in the wild^64^. The *Wolbachia*-positive mosquitoes must reach a sometimes unstable equilibrium level to reliably spread^85^, which could be altered by fitness impacts induced through fungal infection. Further, many agricultural fungal diseases are vectored by arthropods and Wolbachia could be used as a tool to combat disease spread. To begin filling this gap, we sought here to test *Wolbachia*-fungus interactions by systemically infecting the model host *Drosophila melanogaster* with a panel of fungal pathogens and measuring host longevity. We included several variables that we hypothesized might be important factors in any potential pathogen-blocking phenotype, including host genotype, host sex, and pathogen species. We then tested the effect of *Wolbachia* on host fertility and pathogen load when infected or not with fungus.

The main conclusions that can be drawn from the results are that the *w*Mel strain of *D. melanogaster* has a broad, but variable ability to inhibit fungal pathogenesis and that both host and pathogen variables significantly contribute to infection outcomes. Across the systemic infection assays (Figures 1-3, S1-S3), we found a variety of patterns in the results. There are cases where *Wolbachia*-positive flies live significantly longer with fungal infection in all tested contexts, such as *B. bassiana* (Figures 2a, S2a). Notably, this is in agreement with one prior study that showed *D. melanogaster* females with *Wolbachia* lived longer when dipped in a suspension of the same pathogen^72^, suggesting that the phenotype may hold with multiple different infection routes as well. There were also cases where *Wolbachia* significantly increased host longevity in only one host background, such as the *Aspergillus* and *Fusarium* pathogens (Figures 1, S1), *C. rosea* (Figures 2c, S2c), and *Candida* pathogens (Figures 3, S3), examples for which *Wolbachia* was only significant in the *w^k^* background. In contrast, *Wolbachia* was significant in only the *w*^*1118*^ background for *M. anisopliae* infection (Figures 2b, S2b), so either host genotype can result in a statistically significant outcome while the other does not. However, and on a related note, the effect size of *Wolbachia* on host survival may be small in a given context and may lead to lower power to detect the differences with our sample sizes, like *M. anisopliae* in *w^k^* (Figure 2b) or *F. graminaerum* in *w*^*1118*^ (Figure S1d). In contrast, there was one case where the infection outcome was not significant in any context, with the *T. atroviride* pathogen (Figures 2d, S2d), so there may not be an interaction with all pathogens. Further, there were no cases of increased mortality with *Wolbachia*-fungal co-infection, as was suggested in a prior study with fungal pathogens in *Wolbachia*-positive spider mites^73^. Thus, broadly speaking, both pathogen species and host genetics are factors that significantly associate with *Wolbachia*-fungus co-infection outcomes. These patterns suggest that the mechanism(s) of protection are likely not universal to fungal infection, and that host factors are likely involved.

**Figure 3.**
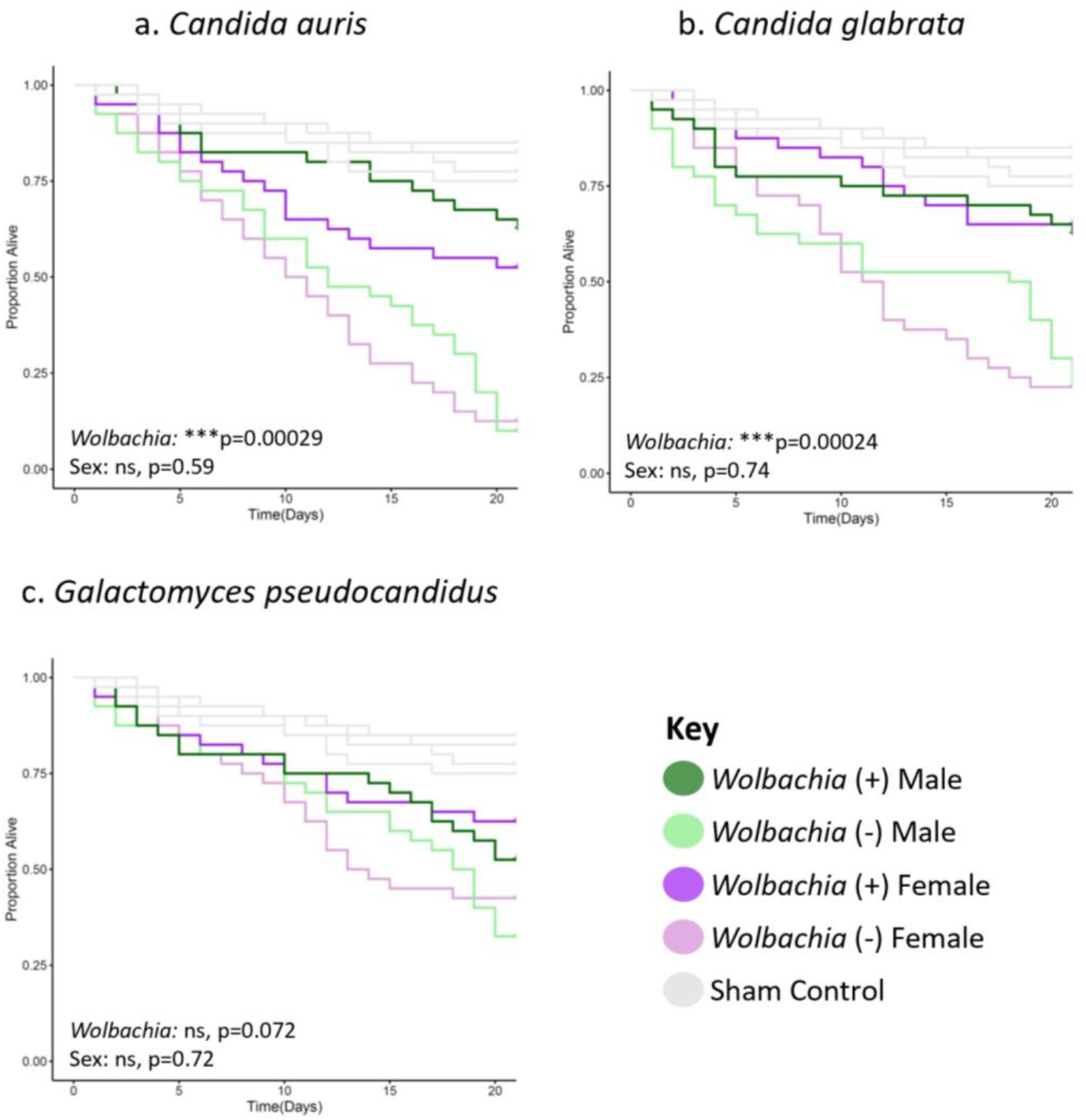
*Wolbachia* increases the longevity of flies of the *w^k^* background line infected with yeast pathogens. Flies of each given background and sex were systemically infected with the indicated pathogen. Infections were performed with either (a) *Candida auris*, (b) *Candida glabrata*, or (c) *Galactomyces pseudocadidus*. Infections of all groups were performed side-by-side, along with those of the *w*^*1118*^ background line (Figure S3), with at least two blocks of infections performed on different days. Each line represents a total of 60 flies. Sham controls were performed with sterile 20% glycerol. Full statistics, available in Table S1, were done with a Cox mixed effects model. Controls are the same in all panels and because they were performed concurrently in the same background.

**Figure 4.**
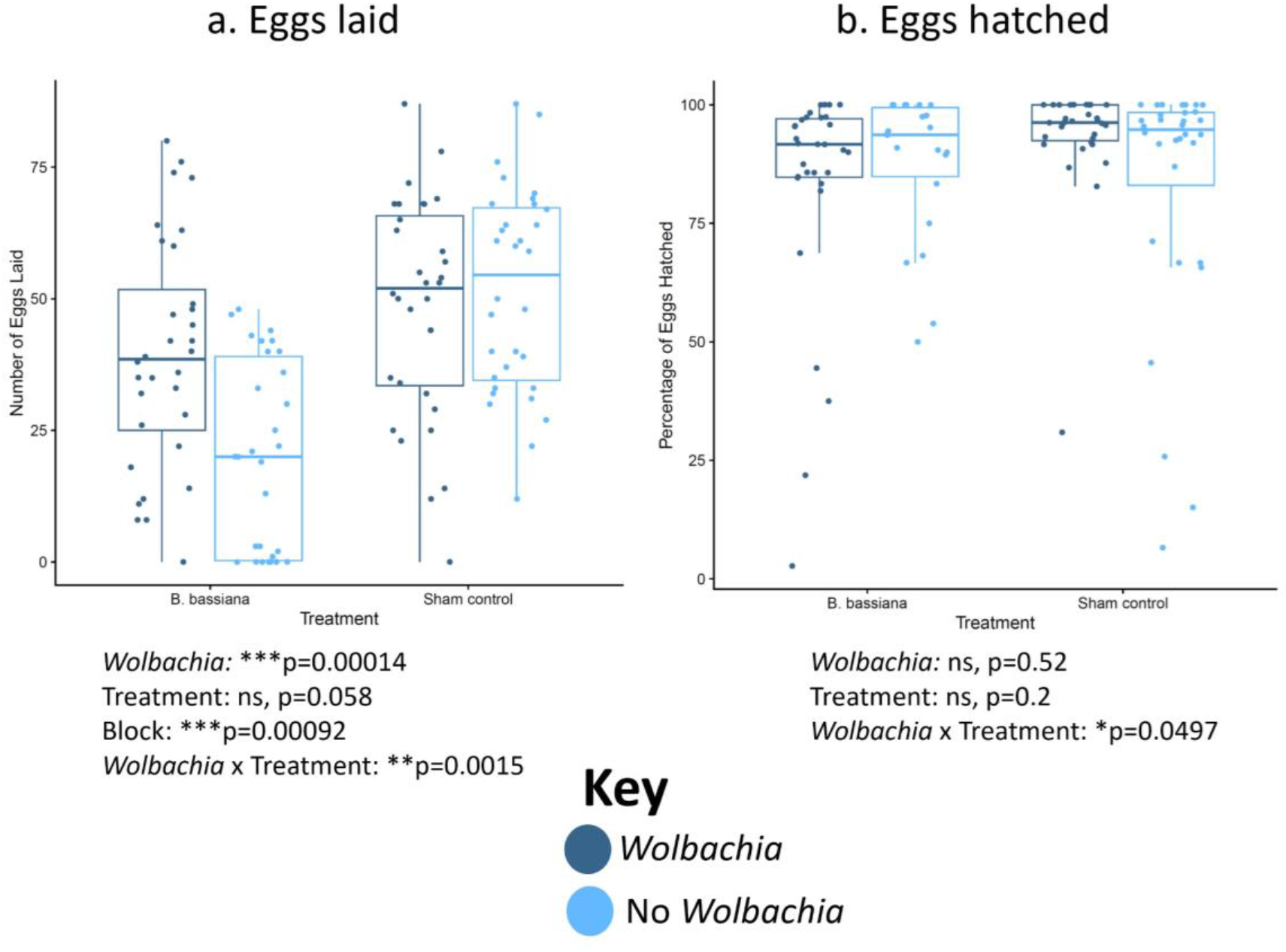
*Wolbachia* increases the number of eggs laid but not the percentage of eggs hatched post-*B. bassiana* infection in the *w^k^* background line. Female flies were systemically infected with *B. bassiana* or treated with a sham control. The flies then laid eggs for 3 days post-infection. (a) Numbers of eggs laid. (b) Proportion of eggs hatched. Each dot represents the total offspring of a single female, with an overall mean of 35 eggs laid. The boxes indicate the interquartile range. Outer edges of the box indicate 25^th^ (lower) and 75^th^ (upper) percentiles and the middle line indicates 50^th^ percentile (median). Whiskers represent maximum and minimum ranges of data within 1.5 times the interquartile range of the box. Statistics are based on a logistic regression (Table S1). The entire experiment was performed twice, and graphs represent a combination of data from both blocks.

**Figure 5.**
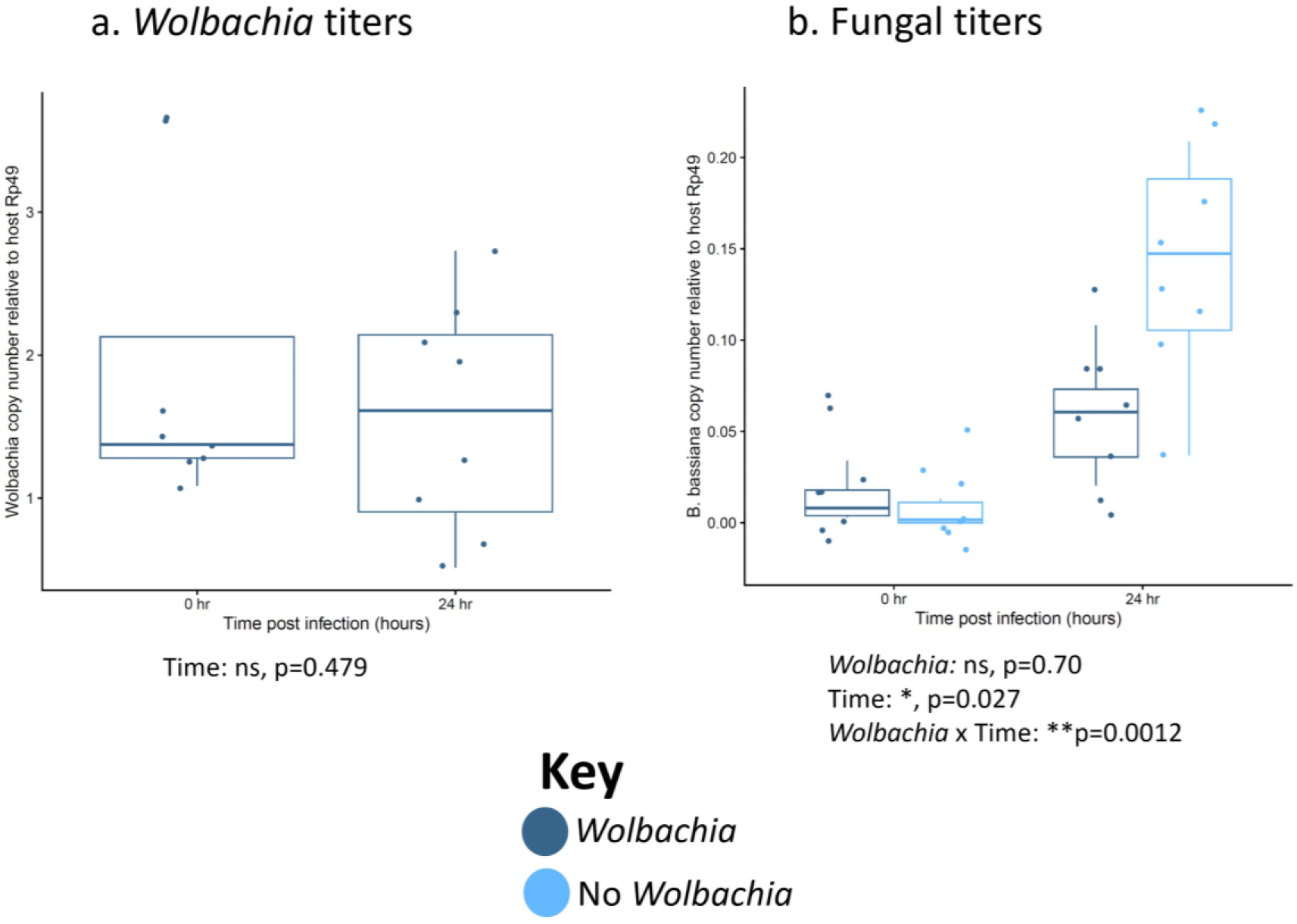
*Wolbachia* associates with reduced pathogen titer after infection with no significant change in *Wolbachia* titer in *w^k^*flies. Female flies were systemically infected with the indicated fungal pathogen and pathogen titers were measured both immediately after infection and 24 h post-infection. Dots represent pools of 3 infected females. (a) *Wolbachia* titers. (b) *B. bassiana* titers. The boxes indicate the interquartile range. Outer edges of the box indicate 25^th^ (lower) and 75^th^ (upper) percentiles and the middle line indicates 50^th^ percentile (median). Whiskers represent maximum and minimum ranges of data within 1.5 times the interquartile range of the box. Statistics are based on a logistic regression (Table S1). The entire experiment was performed twice, and graphs represent a combination of data from both blocks.

Notably, host sex was a significant predictor of infection outcome in several cases as a standalone variable. For example, females had increased longevity compared to males with *B. bassiana* and *M. anisopliae* infection in *w^k^* hosts (Figures 2a,b) and *M. anisopliae* infection in *w*^*1118*^ hosts (Figure S2b), regardless of *Wolbachia* status. In one case, however, male *w*^*1118*^ flies survived at higher rates than females for *T. atroviride* infection (Figure S2d), so the pattern of higher female survival is not always true. Broadly speaking, sex differences in infection outcomes have long been noted in the literature, and are conserved across diverse host and pathogen species^86–88^. Some of the results presented here are also in line with observations that males of many species are often more susceptible to infection than females^89^. Within *Drosophila*, prior research has shown sex differences in infection are common, can favor either males or females, and depend on many different factors^90^. Indeed, infectious challenge with a broad spectrum of bacterial pathogens in *D. melanogaster* demonstrated that females were more broadly susceptible to infection^91^, while another study showed greater female survival with *E. coli* challenge^92^. Those studies identified specific regulators or sensors in both the IMD and Toll pathways that are sexually dimorphic in their expression or activation, contributing to differential immune responses. Sex differences in gut pathology^93^, sexual antagonism in immune resistance and tolerance mechanisms^94^, sex chromosome regulation of immune responses^95^, and sex differences in behavior symptoms^96^ have all been reported for bacterial or viral infections in *Drosophila*. Reports on sex differences in fungal infection have shown mixed results. Notably, several studies have examined sex-specific outcomes of *B. bassiana* infection in *D. melanogaster*. One study showed no sex differences in *D. melanogaster* cuticle infection with *B. bassiana*^97^, another showed higher male survival with *B. bassiana* cuticle infection^98^, and a third also showed higher male survival with *B. bassiana* infection introduced either by spray method or injection^99^. In the third case, removal of various Toll and Imd genes ablated the dimorphism, indicating their role in the phenotype^99^. Notably, the results herein differed, with females showing marginally higher survival with *B. bassiana* infection in the *w^k^* line (Figure 2a), and no sex differences in the *w*^*1118*^ line (Figure S2a). This could be due to differences in the host genetic background strains used in this vs other studies in addition to differences in pathogen infection method or pathogen strain. Thus, sex differences in infection, favoring males or females, are common and the result of many different factors. The fact that we observe sex differences in our results here, but to different extents and in different directions in various contexts, is largely in line with the literature. Future work will be needed to determine basis of these sex differences.

Sex was not only significant predictor of host outcomes alone, but also in combination with *Wolbachia* presence or absence. One particularly interesting case was the significant *Wolbachia* x sex interaction with *F. oxysporum* infection in the *w*^*1118*^ background (Figure S1c). In this case, only *Wolbachia*-positive males survived significantly longer with fungal infection, not females. A similar trend was seen in the *w^k^*background, where statistical significance was evident only when specifically testing within males (Figure 1c). The interaction term of *Wolbachia* x sex was not significant, but these sorts of interactions also suffer from low power. Thus, the mechanism of *Wolbachia* protection from fungal pathogenesis may partially depend on host factors that differ between the sexes, at least in *F. oxysporum* infection. As for why *Wolbachia* may protect males despite transmission mainly through females, it may be due to the dependency of the symbiont on males to induce reproductive parasitism in this species^100^. Notably, the literature investigating *Wolbachia* blocking of viruses and bacteria in arthropods often focuses on one specific sex as opposed to both together, particularly for mosquito research, where viruses are transmitted through female bloodmeals^54,56,101–106^. However, at least one study reports that female *D. melanogaster* infections with Drosophila C Virus are similar to males^51^. Due to few studies comparing the sexes, it is unclear if there are sexually dimorphic outcomes in other cases of *Wolbachia* pathogen blocking or what the molecular and genetic bases of putative *Wolbachia* x sex interactions may be. However, some possibilities include sex differences in *Wolbachia* density, tissue tropism, or dependency on sexually dimorphic host immune responses to inhibit pathogenesis. Future research will be required to investigate this more fully.

Additionally, there was variation in the size of survival differences between *Wolbachia*-positive and -negative flies. In some cases, the difference was small but significant, as with *B. bassiana* (Figures 2a, S2a). In others, the difference was large, such as the *Candida* infections in the *w^k^*background, (Figures 3a,b). Further, there were differences in longevity based on host genetic background, with the *w^k^* flies often succumbing to death earlier, or with fewer overall survivor by the end of the trial period. These results indicate that *Wolbachia*’s impact on fly survival during fungal infection can have a wide range, from only a slight increase in longevity to a much larger one, and that host genetics alone (both sex and genetic background) still significantly influence infection outcomes regardless of *Wolbachia* status. However, even with a modest increase in longevity of a few days for *B. bassiana*-infected flies with *Wolbachia* as an example, the fitness benefits in early stages of infection are significant too (Figures 4, S4). Indeed, the observed increase in early fertility is likely due to reduced pathogen load during initial infection (Figures 4, S4, 5b, S5b). Notably, the lower fungal titers are not due to fluctuating *Wolbachia* titers, as they remain the same during infection (Figures 5a, S5a). This indicates that the symbiont would likely confer a high fitness benefit to a host infected with fungus in the wild due to the combined effects of laying more eggs per day and living more days.

The potential mechanism of fungal pathogen blocking will be the subject of future study. From the reduced pathogen load, it is likely to be an immune resistance mechanism as opposed to tolerance, either of which are known in flies^84,94,107^. In addition, since factors like host sex and genetic background are significant variables, this suggests that the mechanism is likely at least partially mediated through the host. Importantly, the *Wolbachia* strains from each background are nearly genetically identical, with only one single identifiable SNP segregating between the two strains. Although this does not rule out the possibility of differences due to factors like different tissue tropism or DNA structural differences not uncovered by Illumina sequencing, it suggests that differences in phenotypes are likely due to the host rather than symbiont. They do appear to have similar whole-body titers (Figures 5a, S5a), so overall titer probably does not explain any differences. However, future research will need to investigate the relative roles of host and symbiont further. Notably, there is likely to be some overlap in the mechanism(s) of viral and fungal pathogen blocking in *Drosophila*. First, *w*Mel can block both types of pathogens based on the results here and shown elsewhere^51,54,72^. Second, some of the molecular mechanisms contributing to viral blocking could also ostensibly apply to fungal pathogens, such as immune priming^108^, increased ROS production^109^, or competition for resources between symbiont and pathogen^110–112^.

Based on the results, we draw several main conclusions: 1) *w*Mel can confer broad, but not universal, protection against fungal pathogenesis, 2) fungal pathogen blocking by *Wolbachia* is highly context-dependent, with host sex, genetics, and pathogen species being significant determinants of host outcomes, and 3) inhibition of fungal pathogenesis can have positive fitness impacts on the host from early during infection, likely due to reduced pathogen load. Many questions remain unanswered and future work will be needed to investigate this further. For example: How broad is the phenotype in terms of symbiont strains, fly species and strains, and pathogen species? How do other host variables like age impact the phenotype? How do symbiont density and tissue tropism impact the phenotype? Are the results applicable to other insect species for potential translational use in agriculture or other fields? What is the mechanism of fungal pathogen blocking, and can it help inform the mechanism of viral pathogen blocking? How prevalent is fungal pathogen blocking in the wild? This and prior studies pave the way to answering these and other important questions.

## Materials and Methods

### Fly strains and husbandry

Fly strains include *Drosophila melanogaster w*^*1118*^ (one strain with *Wolbachia*, one cured of *Wolbachia* via tetracycline) and *D. melanogaster w*^k^ (one strain with *Wolbachia*, one cured of *Wolbachia* via tetracycline). The *w*^k^ line was isolated in Karsnäs, Sweden in 1960 (*white* allele named for location of isolation)^77^ and the *w*^*1118*^ line was isolated in California and described in 1985 (*white* allele named for date of isolation)^76^. Both were maintained in various labs since their isolation. Flies were reared on CMY media: 64.3 g/L <u>c</u>ornmeal (Flystuff Genesee Scientific, San Diego CA), 79.7 mL/L <u>m</u>olasses (Flystuff Genesee Scientific), 35.9 g/L <u>y</u>east (Genesee Scientific inactive dry yeast nutritional flakes), 8 g/L agar (Flystuff Genesee Scientific *Drosophila* type II agar), 15.4 mL of antimicrobial mixture [50 mL phosphoric acid (Thermo Fisher, Waltham MA), 418 mL propionic acid (Thermo Fisher), 532 mL deionized water], and 1g/L tegosept (Genesee Scientific). Flies were kept at 25°C on a 16h light/8 h dark light cycle.

### Microbial strains and growth conditions for fly infections

The microorganisms used in this study are summarized in Table S2.

Yeast colonies were grown for 16 h on potato dextrose (PD) agar at 30°C. To grow cultures for fly infections, yeast isolates were grown overnight for 16 h from a single colony in 2 mL PD broth (BD, Sparks MA) with shaking at 225 rpm. Isolates were then prepared as described below. Filamentous fungi were prepared by purifying conidia grown on PD agar at 30°C (*Fusarium*, *Aspergillus*, and *Beauveria*) or 25°C (*Metarhizium*, *Clonostachys*, and *Trichoderma*) for 1-2 weeks. Autoclaved DI water was poured over each plate and the conidia were suspended in the liquid. This was then poured over a filter (Millipore Sigma, Burlington MA, Miracloth 22-25 µm pore size) and the filtrate was placed into a 50 mL falcon tube. This was then centrifuged at 1000 rpm for 5 min and the supernatant was discarded. The conidia were then resuspended in sterile 20% glycerol and were counted using a hemocytometer. The conidia concentrations used in this study were (conidia/mL): *Aspergillus fumigatus* (1.75×10^9^), *Aspergillus flavus* (1.18×10^8^), *Fusarium oxysporum* (9.65×10^7^), *Fusarium graminaerum* (1.24×10^8^), *Beauveria bassiana* (4.38×10^8^), *Metarhizium anisopliae* (1.5×10^7^), *Clonostachys rosea* (1×10^8^), and *Trichoderma atroviride* (7.2×10^7^).

### Fly infections

Yeast cultures were grown overnight in the conditions described above. Yeasts *C. glabrata*, *C. auris*, and *G. pseudocandidus* were diluted in PD broth to an optical density (OD) value of A_600_= 200 +/- 5 for *Candida auris* and *Galactomyces pseudocandidus*, and an OD value of A_600_= 220 +/- 5 for *Candida glabrata*. Filamentous fungi were prepared as described above. Mated males or females 4-6 days old of a given genotype were pierced in the thorax just beneath the wing using a 0.15 mm dissecting pin (Entosphinx, Czech Republic, No. 15 Minuten pins 12 mm long 0.15 mm diameter) dipped into the diluted culture or control. Controls were the growth broth for yeasts (PD broth) or sterile 20% glycerol for the filamentous fungi. Flies were then placed in groups of 10 per food vial. 20-30 individuals of each treatment x sex x genotype group were infected in each block, and at least two blocks of infections were performed on separate days for every experiment. Flies were counted for survival daily for 21 days.

### Fertility assay

To measure fertility post-infection, 32 virgin 3-5 day old females were collected from each fly strain (*w*^*1118*^ and *w*^k^, with or without *Wolbachia*). Half of the samples of each strain was infected with *B. bassiana*, as described above. The other half was given 20% glycerol control treatments, also as described above. They were then immediately crossed to 2-4 day old males of the same genotype. Eggs were collected by placing single male-female pairs into a 6 oz. square bottom *Drosophila* bottle (Fisher Scientific, Hampton NH) covered with a grape juice agar plate [100% concord grape juice (Welch’s, MA), tegosept (Genesee Scientific, San Diego CA), 200-proof ethanol (Decon Laboratories Inc, PA), agar (Teknova, Hollister CA), DI water] with yeast paste (Fleischmann’s Active Dry Yeast, Heilsbronn Germany, mixed 1:1 volume with water). These bottles were placed at 25°C incubator overnight. Grape plates were swapped the next morning (16 hr later) with fresh plates and yeast. The bottles were placed back in the incubator and flies were allowed to lay eggs for 72 h. Plates were then removed and eggs were counted immediately. Plates were then kept covered for 24 h and egg hatching was recorded.

### DNA Extractions

DNA extractions were performed with a modified protocol using reagents from the Qiagen Puregene Cell Core Kit (cat. #158046). Cells from samples were lysed by adding 100 µL chilled Cell Lysis Solution to each tube, homogenizing the sample with a pestle, incubating at 65°C for 15 min, then cooling on ice. To precipitate protein, 33 µL Protein Precipitation Solution was added to each sample followed by vortexing for 10 s. Samples were cooled on ice for 5 minutes, and then centrifuged at 14,000 rpm for 3 min. To precipitate DNA, the supernatant was removed and mixed with 100 µL pure isopropanol per sample and each sample was inverted 50 times to mix. The samples were centrifuged 5 min at 14,000 rpm, and supernatant was discarded. Then, 100 µL 70% ethanol was added to each sample and tubes were inverted several times to wash the DNA pellet. Samples were centrifuged 1 min at 14,000 rpm and supernatant was discarded. Tubes were inverted over a paper towel for 10 minutes to dry. DNA was then resuspended with 30 µL DNA Hydration Solution per sample, left at room temperature overnight to allow resuspension, and then frozen and kept at −20°C the next day until use.

### *Wolbachia* and fungal titers

To measure microbial titers post-infection, virgin 3-5 day old females were collected from each fly strain. Flies were then given the indicated treatment, either *B. bassiana* or 20% glycerol sham control. They were then collected at 0 and 24 hr post infection. Samples were flash frozen at their given time point. This led to 10 samples of 3 flies per treatment x time group. This was done for each of the four fly strains.

qPCR was then performed using the Bio-Rad SsoAdvanced Universal SYBR Green Supermix (cat. #1725270) according to manufacturer instructions. Primers are listed in Table S3. qPCR was then performed using a Bio-Rad CFX Connect System with the following conditions: 50°C 10 min, 95°C 5 min, 40x (95°C 10 s, 55°C 30 s), 95°C 30 s. Differences in gene expression were done by calculating 2^-Δct^.

### *Drosophila* and *Wolbachia* sequencing and analysis

For the comparison of the *Wolbachia* from the *w*^*1118*^ and *w*^k^ strains, DNA from 3 female flies each of each strain with *Wolbachia* was extracted as described above. Samples were prepared for whole genome sequencing with the xGen™ DNA Library Prep EZ Kit (Integrated DNA Technologies, #10009821) with a protocol modified to 1/4 reaction volumes. Briefly, 100 ng of DNA from each sample was buffer exchanged via Ampure XP bead purification (Beckman Coulter Life Sciences product number A63881) into the low EDTA TE buffer needed for the xGen™ kit, resulting in a starting input volume of 5 μL. Genomic DNA was enzymatically fragmented to an expected 350 bp insert size, end repaired, and A-tailed in one reaction step. Stubby Y adapters were then ligated onto the fragmented DNA, and reactions were bead-purified following adapter ligation. Unique dual indexes were added to each sample with eight cycles of PCR amplification of the program provided in the xGen™ DNA Library Prep EZ Kit protocol. The libraries were then bead-purified twice, first by a 0.6X purification ratio, followed by a 1.2X purification ratio to provide adapter and primer dimer free libraries. Library quantity was determined with the broad range dsDNA Qubit Assay on the Qubit 1 Fluorometer (Thermofisher Scientific), and the library quality and median library size was assessed with a D1000 screen tape on the TapeStation 4150 (Agilent Technologies). Nanomolar concentrations were determined for each library based on their Qubit concentration in ng/μL and an averaged 442 bp library size. Libraries were pooled at 3 nM concentration along with another set of libraries for a different project. The libraries were sequenced at the University of Kansas Medical Center Genome Sequencing Facility on a NovaSeq 6000 S2 150PE flowcell (Illumina Technologies).

Raw reads were trimmed and filtered using fastp^113^ with default parameters and removing the first and last 5 bases from each sequence. Reads were then mapped to a chimeric assembly of *D. melanogaster* (Release 6 plus ISO1 MT from NCBI) and *w*Mel *Wolbachia* (ASM1658442v1 from NCBI) using bwa^114^ and samtools^115^ with default parameters. SNPs were called using Freebayes^116^ with ploidy set to 1 since the host was inbred and *Wolbachia* is haploid, and filtered with vcffilter^117^ with depth greater than 10 and quality greater than 30.

### Data visualization and statistical analyses

Data analysis and figure generation were performed in R^118^ version 4.2.2, using several packages: coxme^119^ (version 2.2.18.1), ggplot2^120^ (version 3.4.0), cowplot^121^ (version 1.1.1), car (version 3.1.1)^122^, SurvMiner^123^ (version 0.4.9), and SurvMisc^124^ (version 0.5.6). Dot plots were analyzed with a logistic regression. Longevity plots with infection were analyzed using a Cox proportional hazard model with no *Wolbachia* as the reference.

## Data Availability

All data will be deposited in Dryad upon publication of this manuscript.

## Acknowledgments

We would like to thank P. Shahrestani and K. Michel for providing certain microbial strains, as well as J. Blumenstiel for providing fly lines. This work was supported by two National Institutes of Health (NIH) K-INBRE P20 GM103418 postdoctoral awards (to JIP), National Science Foundation (NSF) Postdoctoral Fellowship in Biology (PRFB) DBI 2109772 to JIP, NIH K-INBRE P20 GM103418 student award to AA, and NIH grant R01 AI139154 to RLU.

## Contributions

JIP and RLU conceived, designed, and analyzed experiments and wrote the manuscript. JIP and AA performed fly experiments. MES performed DNA sequencing. All authors approved of the final version of the manuscript.

---

**Figure S1.**
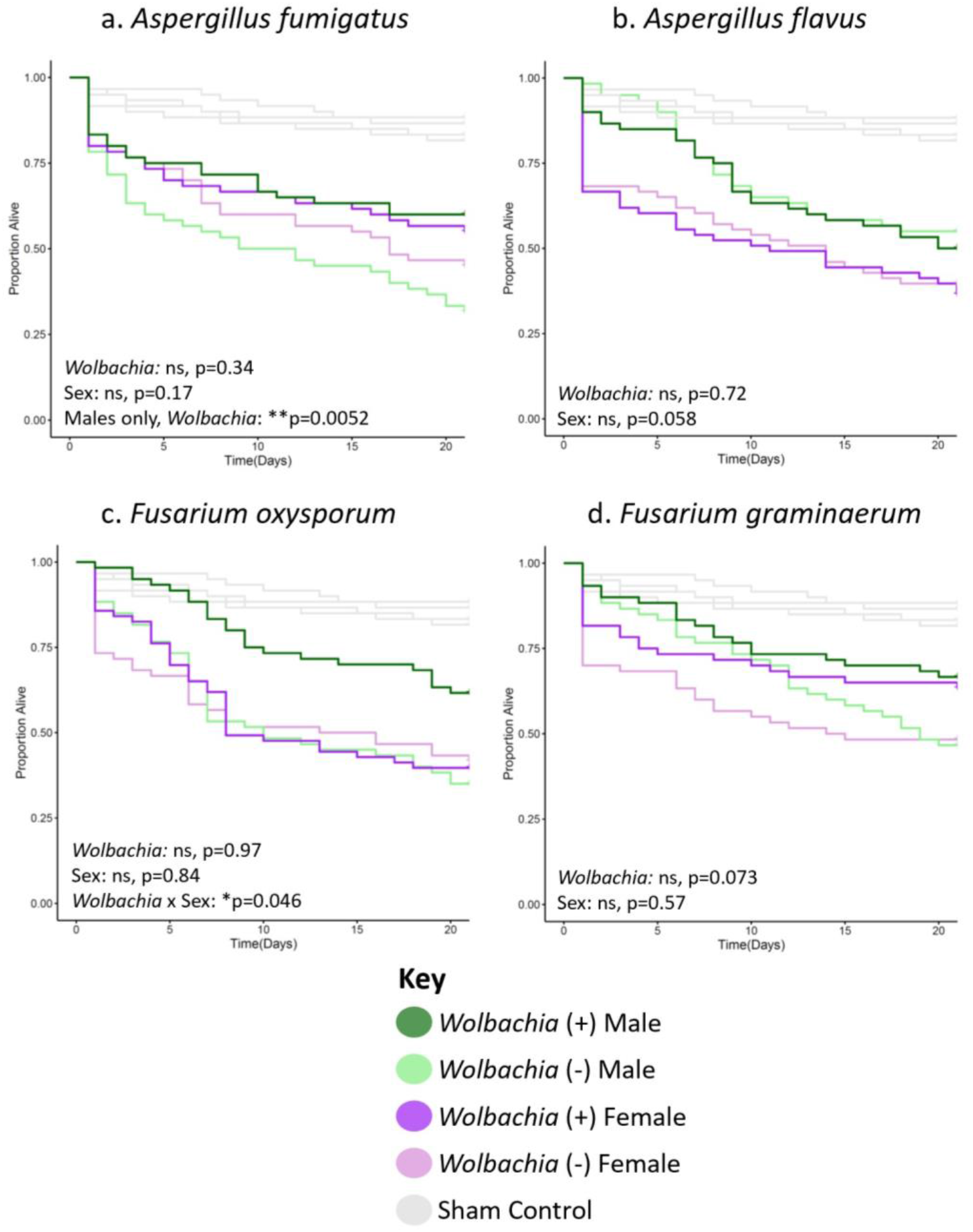
*Wolbachia* does not increase the longevity of flies of the *w*^*1118*^ background line infected with several filamentous fungal pathogens. Flies of each given background and sex were systemically infected with the indicated pathogen. Infections were performed with either (a) *Aspergillus fumigatus*, (b) *Aspergillus flavus*, (c) *Fusarium oxysporum*, or (d) *Fusarium graminaerum*. Infections of all groups were performed side-by-side, along with those of the *w^k^* background line (Figure 1), with at least two blocks of infections performed on different days. Each line represents a total of 60 flies. Sham controls were performed with sterile 20% glycerol. Full statistics, available in Table S1, were done with a Cox mixed effects model. Controls are the same in all panels and in panel S2a because they were performed concurrently in the same background.

**Figure S2.**
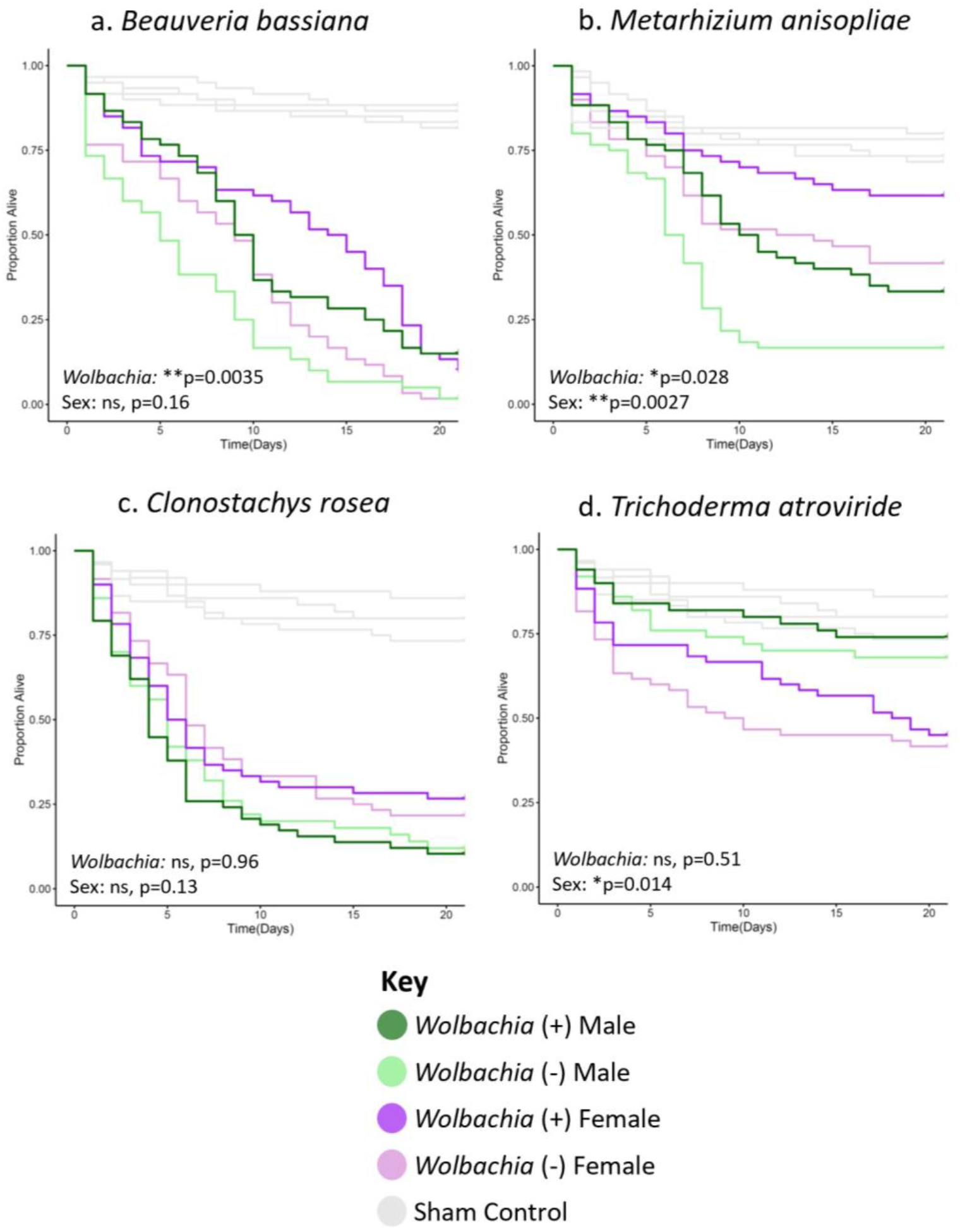
*Wolbachia* increases the longevity of *w*^*1118*^ background line flies infected with certain filamentous fungal entomopathogens. Flies of each given background and sex were systemically infected with the indicated pathogen. Infections were performed with either (a) *Beauveria bassiana*, (b) *Metarhizium anisopliae*, (c) *Clonostachys rosea*, or (d) *Trichoderma atroviride*. Infections of all groups were performed side-by-side, along with those of the *w^k^* background line (Figure 2), with at least two blocks of infections performed on different days. Each line represents a total of 60 flies. Sham controls were performed with sterile 20% glycerol. Full statistics, available in Table S1, were done with a Cox mixed effects model. Controls for panel S2a are the same for Figure S1, and the panels in S2b-d are the same because they were performed concurrently in the same background.

**Figure S3.**
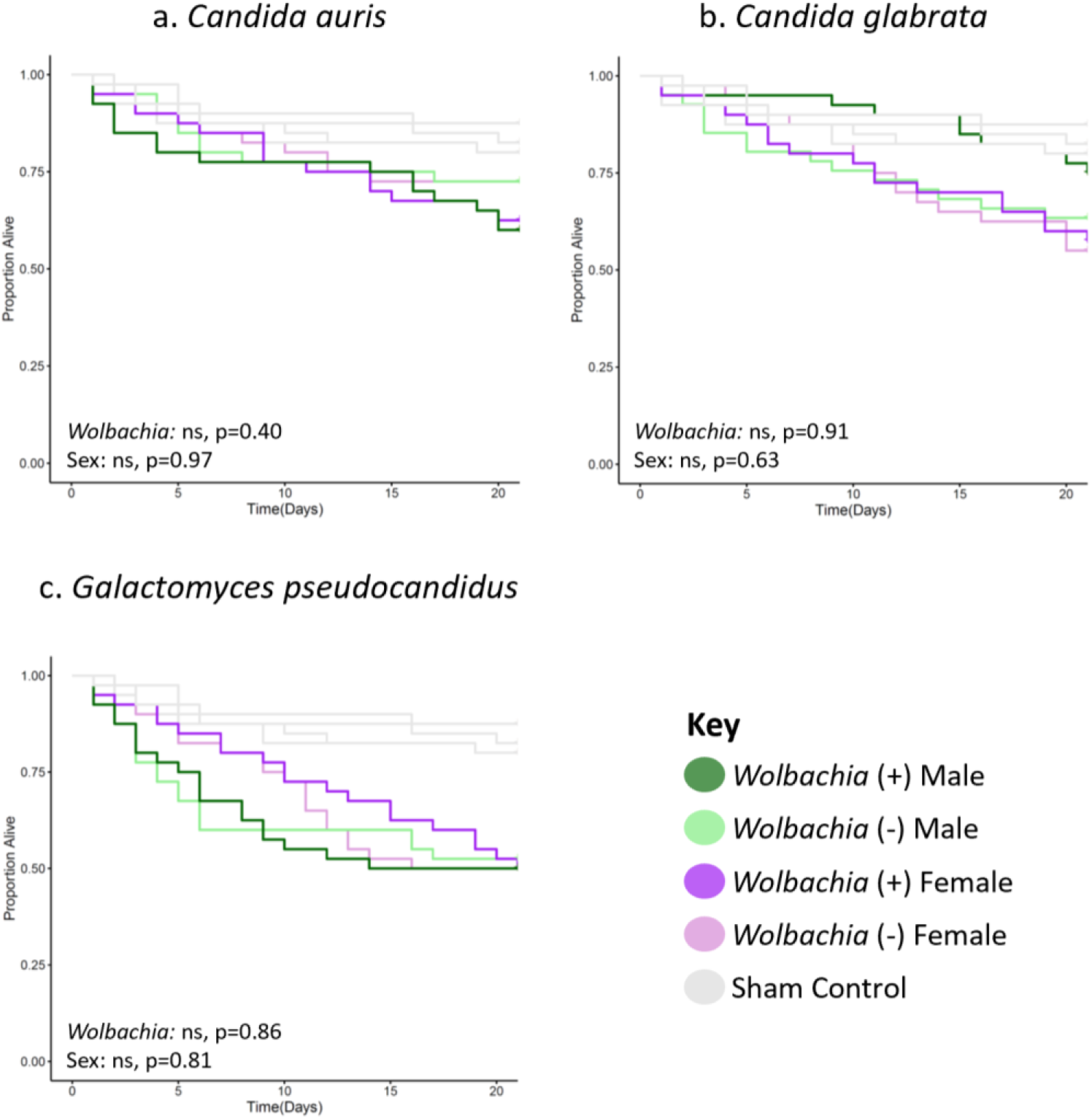
*Wolbachia* increases the longevity of flies of the *w*^*1118*^ background line infected with yeast pathogens. Flies of each given background and sex were systemically infected with the indicated pathogen. Infections were performed with either (a) *Candida auris*, (b) *Candida glabrata*, or (c) *Galactomyces pseudocadidus*. Infections of all groups were performed side-by-side, along with those of the *w*^*1118*^ background line (Figure 3), with at least two blocks of infections performed on different days. Each line represents a total of 60 flies. Sham controls were performed with sterile 20% glycerol. Full statistics, available in Table S1, were done with a Cox mixed effects model. Controls are the same in all panels and because they were performed concurrently in the same background.

**Figure S4.**
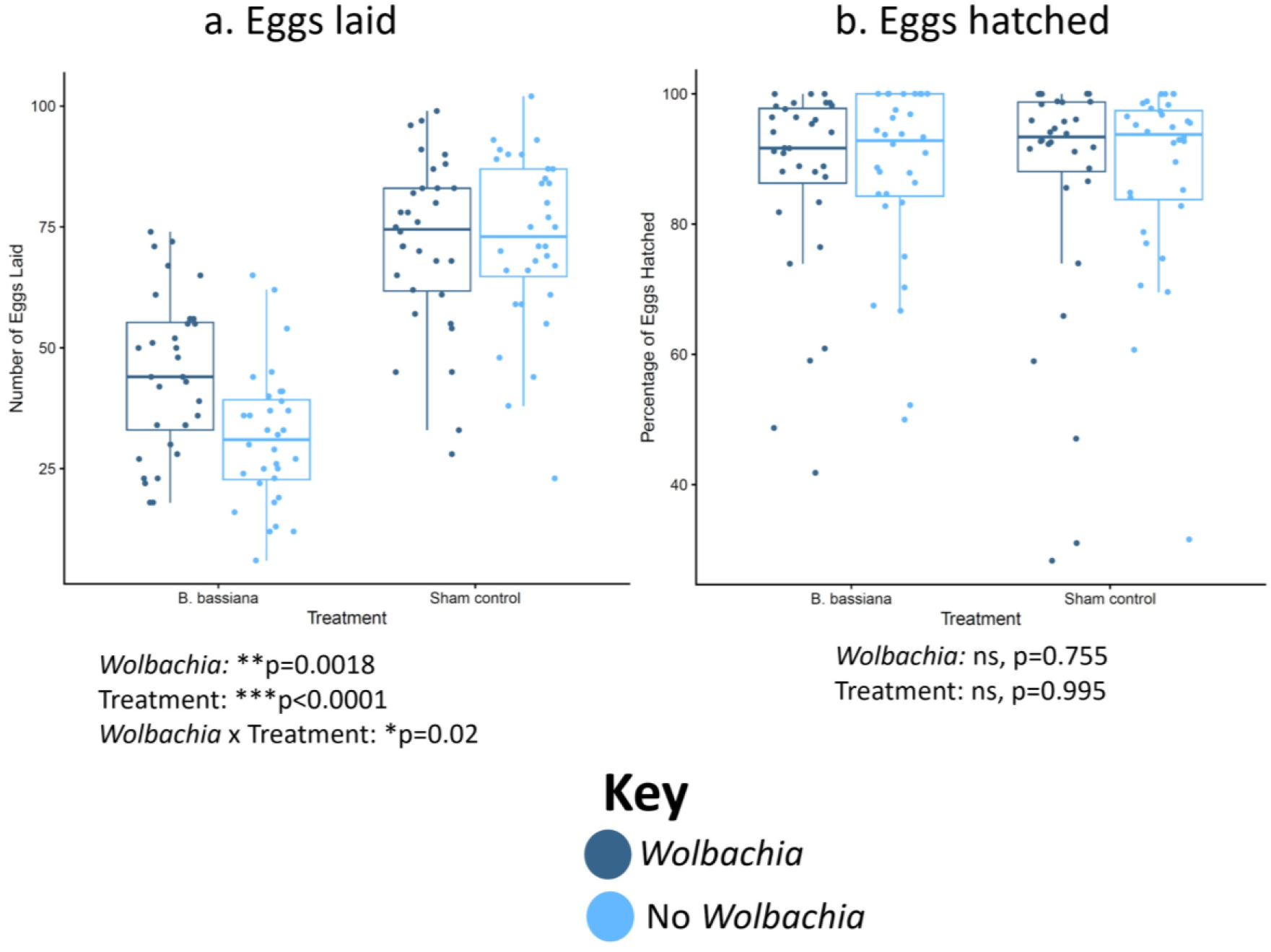
*Wolbachia* increases the number of eggs laid but not the percentage of eggs hatched post-*B. bassiana* infection in the *w*^*1118*^ background line. Female flies were systemically infected with *B. bassiana* or treated with a sham control. The flies then laid eggs for 3 days post-infection. (a) Numbers of eggs laid. (b) Proportion of eggs hatched. Each dot represents the total offspring of a single female, with an overall mean of 48 eggs laid. The boxes indicate the interquartile range. Outer edges of the box indicate 25^th^ (lower) and 75^th^ (upper) percentiles and the middle line indicates 50^th^ percentile (median). Whiskers represent maximum and minimum ranges of data within 1.5 times the interquartile range of the box. Statistics are based on a logistic regression (Table S1). The entire experiment was performed twice, and graphs represent a combination of data from both blocks.

**Figure S5.**
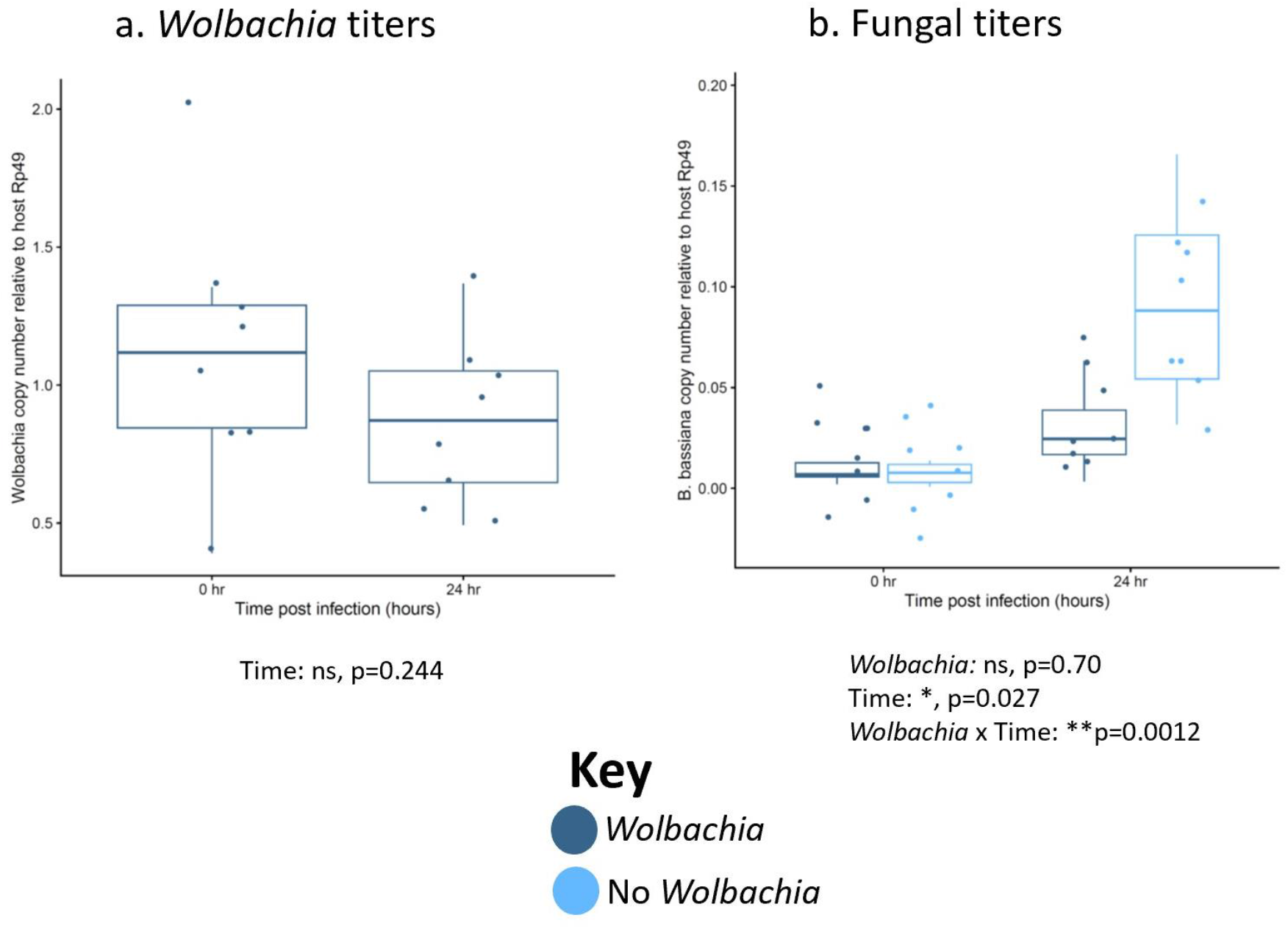
*Wolbachia* associates with reduced pathogen titer after infection with no significant change in *Wolbachia* titer in *w*^*1118*^ flies. Female flies were systemically infected with the indicated fungal pathogen and pathogen titers were measured both immediately after infection and 24 h post-infection. Dots represent pools of 3 infected females. (a) *Wolbachia* titers. (b) *B. bassiana* titers. The boxes indicate the interquartile range. Outer edges of the box indicate 25^th^ (lower) and 75^th^ (upper) percentiles and the middle line indicates 50^th^ percentile (median). Whiskers represent maximum and minimum ranges of data within 1.5 times the interquartile range of the box. Statistics are based on a logistic regression (Table S1). The entire experiment was performed twice, and graphs represent a combination of data from both blocks.

**Table S2.** Microorganisms used in this study.

| Species (strain) | Microbial Classification | Isolation Source | Stock Number or Isolated/Gifted By |
| --- | --- | --- | --- |
| *Candida glabrata* (CBS 138) | Yeast | Feces | ATCC 2001 |
| *Candida auris* | Yeast | Clinical isolate | CDC B11903 |
| *Galactomyces pseudocandidus* | Yeast | *Drosophila* | Isolated by I. Nevarez-Saenz |
| *Fusarium oxysporum* (f. sp. *Lycopersici*) | Filamentous fungus | Tomato | FGSC 9935 |
| *Beauveria bassiana* (GHA) | Filamentous fungus | *Locusta migratoria* | Gift from P. Shahrestani |
| *Aspergillus fumigatus* | Filamentous fungus | Clinical isolate | FGSC 1100 |
| *Aspergillus flavus* (NRRL 3357) | Filamentous fungus | Peanut | FGSC A1446 |
| *Metarhizium anisopliae* (recently renamed *Metarhizium robertsii*) | Filamentous fungus | Insect | ARSEF 23 |
| *Clonostachys rosea* | Filamentous fungus | *Aedes albopictus* (mosquito) L4 larvae, Manhattan, KS | Isolated by P. Tawidian & gifted by K. Michel |
| *Trichoderma viride* | Filamentous fungus | *Aedes albopictus* (mosquito) L4 larvae, Manhattan, KS | Isolated by P. Tawidian & gifted by K. Michel |

**Table S3.**
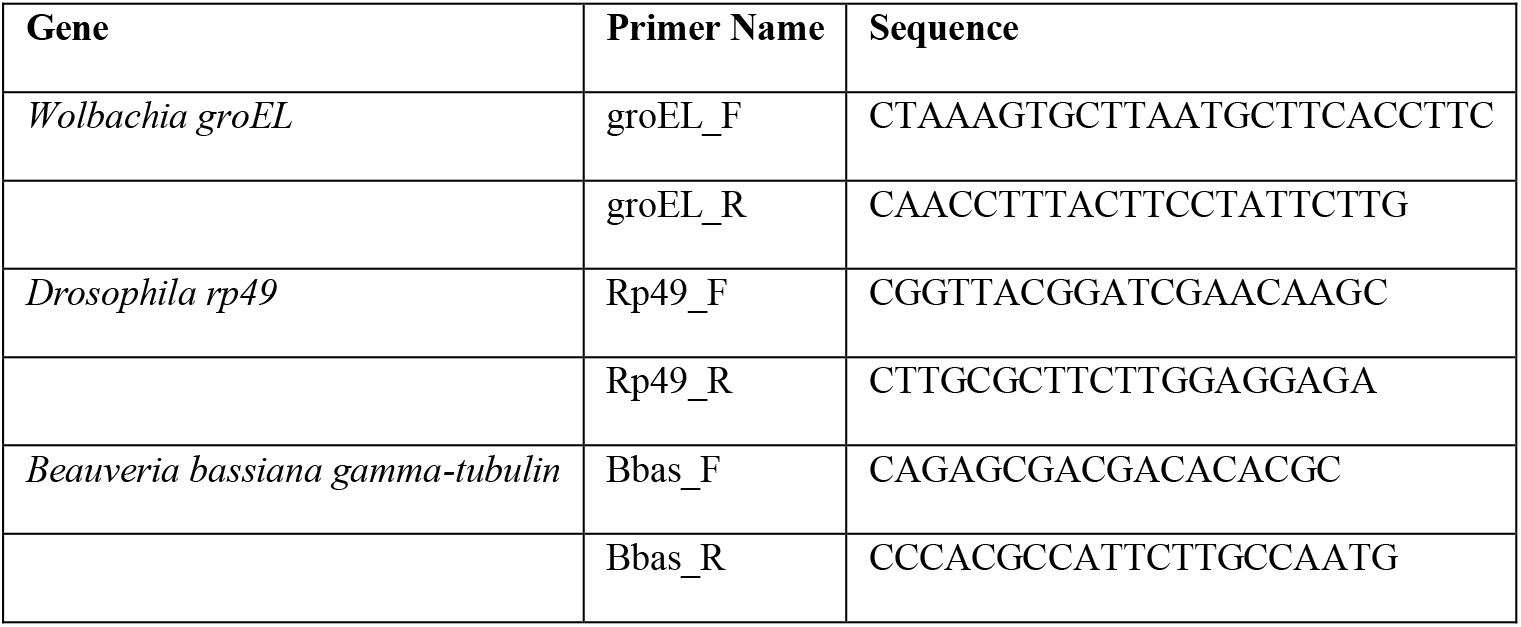
Primers used in this study.

